# Methods for Implementing Integrated Step-Selection Functions with Incomplete Data

**DOI:** 10.1101/2023.11.08.566194

**Authors:** David D. Hofmann, Gabriele Cozzi, John Fieberg

## Abstract

Integrated step-selection analyses (iSSAs) are versatile and powerful frameworks for studying habitat and movement preferences of tracked animals. iSSAs utilize integrated step-selection functions (iSSFs) to model movements in discrete time, and thus, require animal location data that are regularly spaced in time. However, many realworld datasets are incomplete due to tracking devices failing to locate an individual at one or more scheduled times, leading to slight irregularities in the duration between consecutive animal locations. To address this issue, researchers typically only consider bursts of regular data (i.e., sequences of locations that are equally spaced in time), thereby reducing the number of observations used to model movement and habitat selection. We reassess this practice and explore four alternative approaches that account for temporal irregularity resulting from missing data. Using a simulation study, we compare these alternatives to a baseline approach where temporal irregularity is ignored and demonstrate the potential improvements in model performance that can be gained by leveraging these additional data.

## 1 Introduction

Understanding how animals move across the landscape, what habitats they prefer, and what resources they select are fundamental questions in movement ecology (Nathan, 2008). Thanks to recent advances in animal tracking (Cagnacci et al., 2010; Williams et al., 2019; Beardsworth et al., 2022) and remote sensing technologies (Toth & Jóźków, 2016; Rumiano et al., 2020), new opportunities and analytical tools have emerged for studying how animals move and interact with their environment (Tomkiewicz et al., 2010; Kays et al., 2015). Methods commonly used to analyze animal movement data, including step-selection analyses (Fortin et al., 2005; Thurfjell et al., 2014; Fieberg et al., 2021) and hidden Markov models (Michelot et al., 2016), require **animal locations** (terms in bold at first occurrence are defined in Table 1) that are collected at a constant sampling frequency, leading to data that are equally spaced in time. Yet, it is common to encounter missing locations in most telemetry data sets (Frair et al., 2010; Hofman et al., 2019; Vales et al., 2022), which introduces unwanted irregularities in the duration between successive locations. Thus, there is a need for analytical tools that enable the analysis of such data, while mitigating potential biases arising from temporal irregularity introduced through missing animal locations.

**Table 1:** Glossary of terms. Terms in the glossary are printed in bold at first occurrence in the main text. Definitions are always given in the context of step selection functions (SSFs).

| Term | Definition |
| --- | --- |
| **Animal locations** | A series of telemetry data points that include date, time, longitude, and latitude information, describing when and where an animal was observed or recorded. |
| **Step** | A straight line connecting two consecutive locations. |
| **Observed step** | A step that connects two observed animal locations. |
| **Random step** | A step that connects an observed animal location with a random location. Random locations are typically generated by combining an observed animal location with random step-lengths and turning-angles. |
| **Step-length** | The Euclidean distance of a step. |
| **Turning-angle** | A measure of the change in direction between two consecutive steps. |
| **Habitat-selection function** | A probabilistic description of an animal’s habitat preferences. It describes how an animal selects habitat when not constrained by its movement capacity. Also referred to as movement-free habitat selection function. |
| **Movement kernel** | A probabilistic description of an animal’s movement capacity. It describes how an animal would move when not constrained by habitat selection. Also referred to as selection-free movement kernel. |
| **Trajectory** | A sequence of animal locations collected on the same individual. |
| **Step-duration** | The time interval associated with a particular step, i.e., the time elapsed between two consecutive animal locations. |
| **Regular animal locations** | A series of animal locations that have been obtained at regularly-spaced time intervals, such as every hour. |
| **Irregular animal locations** | A series of animal locations collected at irregular time intervals. |
| **Regular step-durations** | Step durations that occur when animal locations are successfully collected at regular time intervals. |
| **Irregular step-durations** | Step durations that occur when animal locations are not successfully collected at regular time intervals. |
| **Missingness** | The fraction of animal locations that should have been collected but, for some reason, were not. For example, if only eight out of ten expected animal locations were successfully collected, the missingness would be 0.2 (i.e., 20%). |
| **Forgiveness** | The maximum step-duration, measured in multiples of the regular step-duration, a modeler is willing to include in the step-selection analysis. A modeler with a forgiveness of one, for instance, only considers regular steps, while a modeler with a forgiveness of two would consider irregular steps up to twice the regular step-duration. |
| **Burst** | A sequence of consecutive animal locations equally spaced in time and with steps were the step-duration does not exceed the forgiveness. |
| **Valid step** | A step for which a step length, turning angle, and step duration can be computed, and for which the step duration does not exceed the forgiveness of the modeler. These steps can be used for step-selection analysis. |

Step-selection analyses (SSAs) are widely used to study animals’ movement capacities and habitat-selection patterns (Fortin et al., 2005; Thurfjell et al., 2014). Straight-line segments connecting consecutive animal locations, referred to as **steps**, form the basic building blocks of the statistical likelihood in SSAs. Specifically, SSAs model the probability *u* of finding an individual at location *s* at time *t* + 1, given the animal’s past positions at time *t* and *t−*1, *s*_*t*_ and *s*_*t−*1_, respectively:

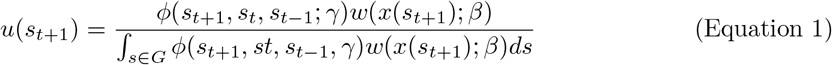

Here, the function *ϕ* represents an animal’s **movement kernel** which is usually expressed in terms of **step-length** and **turning-angle** distributions, with *γ* representing parameters in these distributions. The function *w* is the **habitat-selection function** and reflects an animal’s preferences *β* for environmental characteristics *x* at location *s*_*t*+1_. In most applications, *w* is modeled as a log-linear function of *x*, taking the form *w* = *exp*(*x*^*T*^*β*). The integral in the denominator of Equation 1 ensures that *u* is a proper probability distribution (i.e., that it integrates to 1). Following Michelot et al. (2023), we call the product *ϕ* × *w* the step-selection function (SSF), as it highlights that the probability of finding an animal at a certain location depends on both the animal’s movement kernel and its habitat-selection function.

Given a series of observed steps, finding the movement and habitat-selection parameters that maximize the likelihood in Equation 1 requires approximating the integral in the denominator for each observed step. A variety of numerical integration techniques can be used for this purpose (Michelot et al., 2023), but a common approach is to combine **observed steps** with **random steps** generated by sampling **step-lengths** and **turning-angles** from parametric distributions informed by the data (Fortin et al., 2005; Thurfjell et al., 2014). Environmental conditions at observed steps are then contrasted with environmental conditions at random steps in a (mixed effects) conditional logistic regression framework (Fortin et al., 2005; Muff et al., 2020). To jointly estimate parameters in *ϕ* and *w*, movement descriptors (e.g., step-length (sl), its natural logarithm (log(sl)), and the cosine of the turning-angle (cos(ta))) can be included in the conditional logistic regression model, and their estimated coefficients can be used to update the initial (tentative) step-length and turning-angle distributions (Duchesne et al., 2015; Avgar et al., 2016; Fieberg et al., 2021). The specific descriptors that need to be included depend on the assumed step-length and turning-angle distributions (for more details, see Appendix C of Fieberg et al., 2021). This approach to estimating parameters of the SSF, termed *integrated* SSA (or iSSA) by Avgar et al. (2016), is similar to using importance sampling to approximate the integral in Equation 1 (Michelot et al., 2023) and is readily accessible through the R-package amt (Signer et al., 2019).

SSAs have proven extremely effective in numerous ecological studies (Thurfjell et al., 2014), providing insights into seasonal space use (Vales et al., 2022; Enns et al., 2023), resource selection during distinct behavioral phases (Elliot et al., 2014; Abrahms et al., 2017; Cozzi et al., 2018; Broekhuis et al., 2019), and the effects of landscape familiarity or memory on animal movements (Kim et al., 2023). A model parametrized using iSSA resembles a fully mechanistic movement model that can be used to simulate space use under novel conditions (Avgar et al., 2016; Signer et al., 2017; Hofmann, Cozzi, McNutt, et al., 2023; Signer et al., 2023). This characteristic has made iSSAs a useful tool for quantifying landscape resistance and identifying movement corridors (Buchholtz et al., 2020; Zeller et al., 2020; Hofmann et al., 2021; Hofmann, Cozzi, McNutt, et al., 2023).

A key assumption when conducting an iSSA is that the data have been collected at a constant sampling frequency, thus producing **trajectories** with **regular step-durations** (Δ*t*; Fortin et al., 2005; Thurfjell et al., 2014). Here, we refer to such data as **regular animal locations**, and without loss of generality, we assume the regular step-duration to be one (i.e., Δ*t* = 1). Regular step-durations ensure that step-lengths and turning-angles are independent of the step-duration and therefore steps can be pooled when estimating tentative movement parameters. Since animal locations are usually obtained using automated tracking devices, such as GPS collars programmed to record data at regular intervals, satisfying this assumption seems straightforward. In reality, however, device limitations often imply that some of the aspired datapoints fail to be collected, thus introducing **missingness** and confronting researchers with **irregular animal locations** and **irregular step-durations** (Frair et al., 2010). In a comprehensive study, Hofman et al. (2019) showed that across 3,000 GPS devices and 160 species, the average success rate of obtaining a scheduled animal location was 78% (implying a missingess of 22%), thus highlighting that irregular animal locations are a frequent phenomenon in ecological studies.

It is generally recommended that, in the case of such irregular data, researchers should only retain **bursts** of steps with regular step-durations (possibly with some tolerance) and discard the rest (Thurfjell et al., 2014). We will refer to this modeling approach as having a **forgiveness** level of one, indicating that only steps with step-durations Δ*t* = 1 are retained for further analysis. In R, the `amt` package provides the function `track_resample` specifically for identifying bursts of steps with regular step-durations (Signer et al., 2019). The main drawback of this approach is that it may result in a substantial amount of data being discarded (Figure 1 and Figure 2). For instance, consider a hypothetical trajectory in which location 4 is missing (Figure 1a). The absence of this location prevents the computation of a step between locations 3 and 4, as well as between locations 4 and 5. Furthermore, without these steps, it becomes impossible to compute a turning-angle for the step between locations 5 and 6. Consequently, the lack of a single location reduces the effective sample size, which is the number of **valid steps**, by three. Assuming that animal locations are missing at random, a missingness level of 25% causes the number of valid steps to drop by 58% (Figure 2). A modeler willing to increase their level of forgiveness to two (i.e., allowing for inclusion of steps with Δ*t* ≤ 2) would be able to increase the number of valid steps by 56% (Figure 1 and Figure 2), therefore achieving a substantial gain in effective sample size. The ability to capitalize on irregular data is likely to be particularly important for applications where data are already limited, such as, for instance, when modeling dispersing individuals (Rudnick et al., 2012; Fattebert et al., 2015; Cozzi et al., 2020). However, increasing the forgiveness also implies that step-durations of the retained steps become irregular, thus necessitating appropriate tools to account for such irregularity.

**Figure 1:**
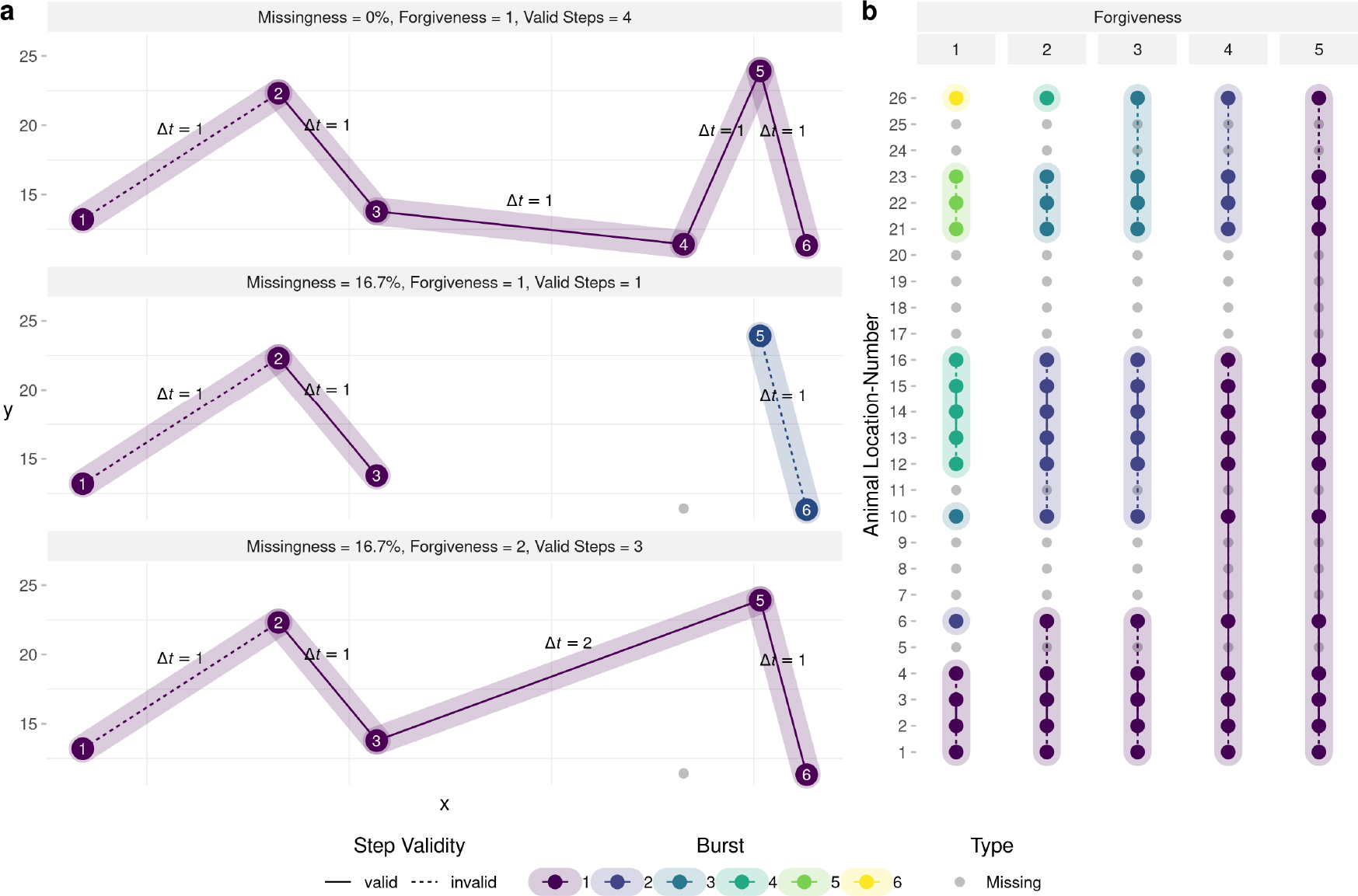
(**a**) Demonstration of how missingness affects the number of valid steps that can be used for step-selection analyses under different levels of forgiveness. The upper panel depicts a trajectory with zero missing locations. That is, all aspired locations were successfully collected on a regular interval (yielding a regular step-duration of Δ*t* = 1). This trajectory produces four valid steps that can be included in the iSSF model and one invalid step that has to be omitted because it has no turning-angle associated with it. In the central panel, animal location 4 was not obtained, introducing a missingness of 16.7%. If the modeler has a forgiveness of one, only a single step can be included for further analysis, as all other steps are invalid (either because no turning-angle can be computed or because step-durations exceed the forgiveness). If, however, the modeler exhibited a forgiveness of two, such as in the lower panel, a total of three steps could be obtained for further analysis. (**b**) Conceptual illustration of how increasing the forgiveness allows one to retain additional steps that can be used for step-selection analysis. The sequence of dots resembles the sequence of locations that were scheduled to be collected (e.g. using a GPS device), with the lines representing hypothetical steps. Because not all locations were successfully obtained (gray dots), there is missingness. Depending on the forgiveness level, already a single missing location enforces the introduction of a new burst, which leads to the loss of several steps. In addition, some of the remaining steps are invalid (dotted) because they are lacking a turning-angle. By increasing the forgiveness, a modeler is willing to retain steps that exceed the regular step-duration by a certain threshold, which enables them to obtain longer bursts and increase the number of steps that can be used for further analysis. In the figure, forgiveness increases from left to right.

**Figure 2:**
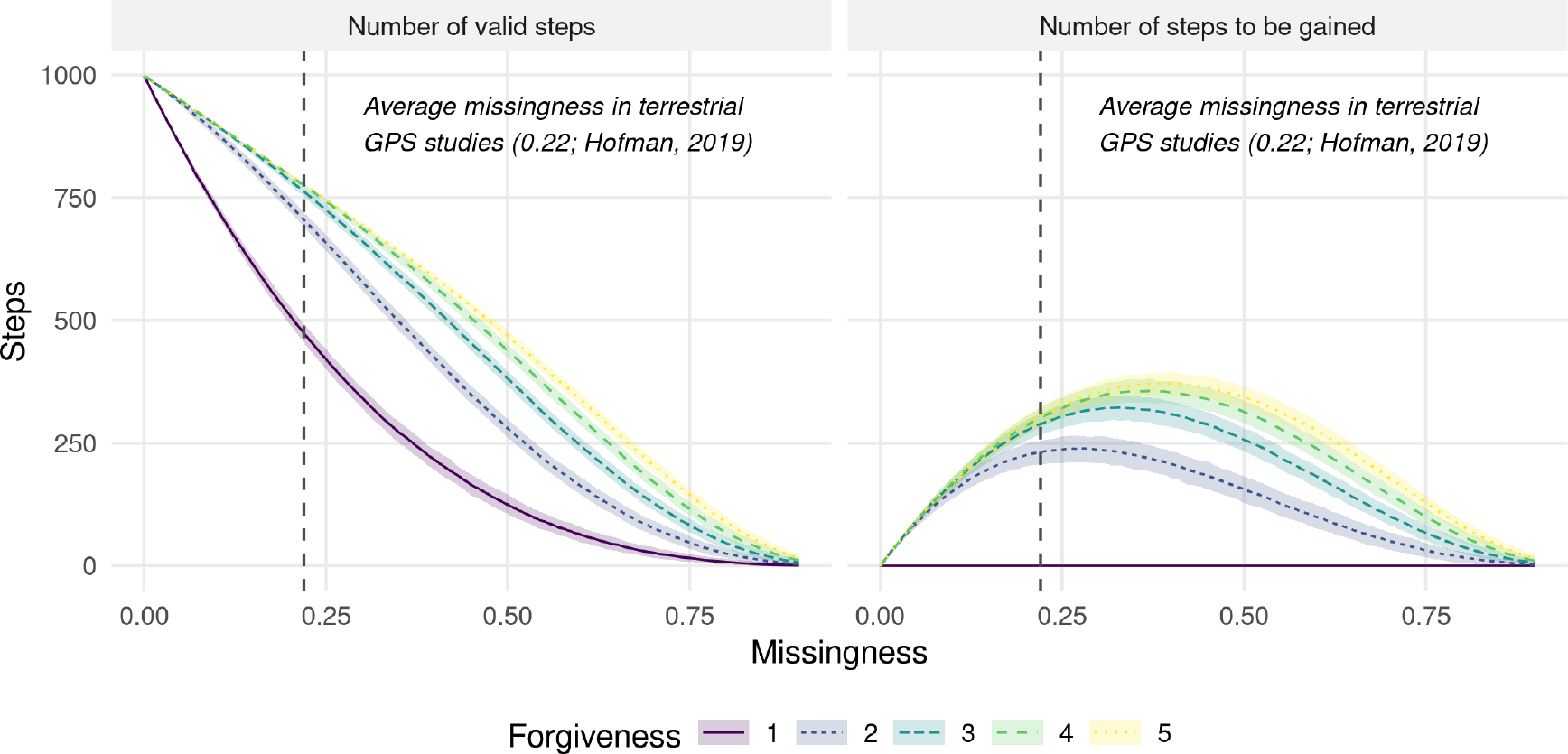
Illustration of how missingness in animal location data reduces the number of valid steps that can be used in step-selection analyses (left panel) and how increasing forgiveness helps to retain additional steps that are otherwise omitted (right panel). At a missingness of 0, 998 valid steps can be obtained from the total of 1,000 animal locations. At higher missingness, step-durations become irregular, which means that the number of valid steps decreases substantially. However, if the modeler is willing to increase their forgiveness, additional steps can be gained. The right panel shows the number of valid steps that can be gained is when increasing the forgiveness from 1 to 2, 3, 4, and 5, respectively. Ribbons indicate the 95%-percentile intervals derived from 1000 replicates.

Various methods have been employed in the past to address temporal irregularities in animal location data and may serve as valuable starting points for developing approaches that enable the integration of irregular data with iSSFs.

### 1. Imputation

An intuitive solution is provided by McClintock (2017), who suggests fitting a continuous-time correlated random walk movement model (Johnson et al., 2008) to the collected data and to use the fitted model to impute missing fixes. By imputing missing locations, the analysed trajectories become entirely regular again and can be analysed using traditional techniques. This approach, which we coin the *imputation* approach, is readily available through the `R`-package `crawl` (Johnson et al., 2022), yet has only been tested for use with hidden Markov movement models and not with iSSFs (McClintock, 2017).

### 2. Naïve

Another approach is outlined by Munden et al. (2021), who introduced timevarying iSSFs. In this framework, a change-point detection algorithm is applied to the series of observed animal locations to identify distinct decision points where the animal turns (Potts et al., 2018; Munden et al., 2021). Steps are then created to represent straight-line movements in-between these decision points, but because decision points are not regularly spaced in time, the resulting step-durations are irregular. Thus, stepdurations are treated as random variables, and, instead of generating random steps by sampling step-lengths and turning-angles, the authors generate random steps by sampling step-durations, step-speeds, and turning-angles. The underlying assumption is that step-lengths scale linearily with step-duration and can therefore be meaningfully represented by combinations of step-speeds and step-durations. Although this approach was developed with ultra-high-frequency data in mind, we might *naïvely* apply it more broadly to the case of missing data if we believe the presumed linearrelationship to hold true (this assumption may be reasonable when step-durations are short, but it is unlikely to hold for longer step-durations since an animal’s path will deviate from a straight line between successive locations). Hence, we propose with our *naïve* approach, to scale the generated random steps by the observed step-duration.

### 3. Dynamic+Model

Instead of generating random steps by sampling step-lengths and turning-angles from distributions fitted to a single step-duration, one may choose to fit separate distributions to steps of different durations, thus acknowledging potentially non-linear relationships between step-duration and step-lengths or turning-angles. Because random steps in this approach are sampled using different *tentative* distributions, it is necessary to include interactions between step-durations and other step-descriptors (sl, log(sl), and cos(ta)) in the conditional logistic regression model to allow updating tentative distributions to the different step-durations. We therefore refer to this approach as the *dynamic+model* approach, highlighting that step-length and turningangle distributions are dynamically adjusted to observed step-durations and that the step-duration is included as a modifier of the coefficients of step-descriptors in the regression model.

### 4. Multistep

Finally, we propose a *multistep* approach, where random steps of varying step-durations are generated by stitching together sequences of random steps from the regular step-duration. For example, one can generate a random step of duration Δ*t* = 2 by combining two random steps of step-duration Δ*t* = 1.

Our goal with this article is to reassess the practice of discarding irregular animal locations in iSSFs and to investigate whether retaining irregular data could, in fact, serve to improve model performance. Our hypothesis is that even irregular data contains valuable information on habitat and movement preferences that could be leveraged if appropriate methods are applied. To test this notion, we conducted a comprehensive simulation study where we simulated regular animal location data with known movement and habitat parameters. We then introduced varying levels of missingness and applied iSSFs to estimate simulation parameters. Specifically, we employed the four alternative iSSF approaches outlined above and compared them to the traditional approach of including only bursts of regular data and to an uncorrected approach that simply ignored irregular step-durations when using a forgiveness level *>* 1. To examine the impact of different landscape configurations on derived estimates, we ran our simulations for different levels of spatial autocorrelation. The use of simulations instead of real data had the benefit that underlying parameters of the movement kernel and habitat-selection function were known, which allowed us to assess the reliability of different methods in retrieving true simulation parameters under different conditions (sensu Kéry and Royle, 2016).

We anticipated that increasing forgiveness without adjusting for the introduced irregularity would entail a bias-variance trade-off. Specifically, we anticipated that increasing forgiveness would allow improving estimator precision, but at the cost of introducing bias due to failure of accounting for irregular sampling intervals. We expected this bias to be particularly pronounced at high levels of missingness. Furthermore, we hypothesized that accounting for irregularity in the *naïve, dynamic+model*, and *multistep* approaches would improve model accuracy, while alleviating potential bias, thus providing an effective means of incorporating additional data. Because the *imputation* approach relied on an intermediate movement model to predict missing animal locations, we had no prior expectations for how well it would perform.

## 2 Methods

We implemented the simulation study in the programming language `R` version 4.3.2 (R Core Team, 2023) and achieved parallelization of simulation-runs using the `R`-package `pbmcapply` (Kuang, 2022). We generated figures using the `ggplot2` (Wickham et al., 2023), `ggpubr` (Kassambara, 2023), and ggh4x (Brand, 2023) `R`-packages. We manipulated `raster` data and computed spatial distances using the R-package raster (Hijmans et al., 2023). An overview of the simulation design and the different iSSF approaches is presented in Figure 3 and all codes to reproduce this study are available through an online repository (Hofmann, Cozzi, & Fieberg, 2023).

**Figure 3:**
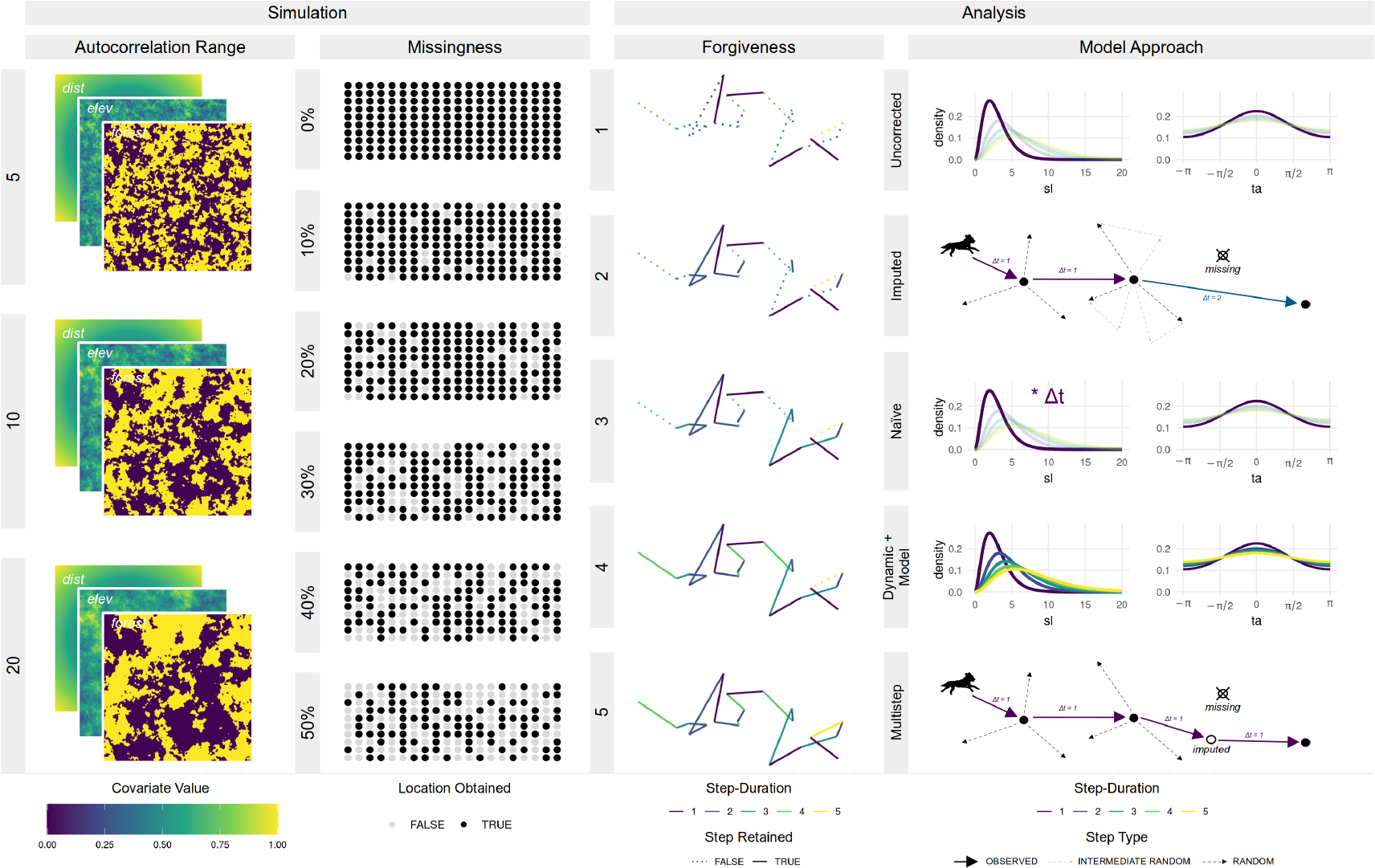
Illustration of the study design. We varied the autocorrelation range when simulating spatial covariates from 5 to 20 and tested for different missingness scenarios (ranging from 0% to 50% missing locations). To investigate how increasing forgiveness (i.e., the willingness to include steps with duration above the regular step-duration) influenced model results, we varied its value from 1 (regular step-selection) to 5 (considering steps that are five times the regular step-duration). Finally, we tested five different methods to account for potential biases introduced by including irregular steps. This gave 3 x 6 x 5 x 5 = 450 combinations, each of which we replicated 100 times. We assumed step-lengths (sl) to follow a gamma distribution, whereas turning-angles (ta) followed a von Mises distribution.

### 2.1 Landscape Simulation

We simulated a virtual landscape comprising two continuous and one categorical (binary) spatial layers, each with a resolution of 300 x 300 pixels (Figure 4) and spanning across xand y-coordinates from 0 to 300. The first layer, `Dist` (continuous), quantified the distance to the center of the virtual landscape (*x* = 150, *y* = 150), and can be understood as a point of attraction, such as, for instance, the center of an animal’s home-range. The second layer, Elev (continuous), resembled an elevation layer and was simulated by randomly sampling values from a normal distribution for each pixel. To achieve spatial autocorrelation, we applied a circular moving window with radius *r* within which we tallied pixel-values. We varied *r* from 5, to 10, to 20, depending on the simulated level of autocorrelation (Figure S2). The third layer, `Forest` (categorical), represented areas covered by woodland and was simulated similarly to the `Elev` layer, but we binarized the layer by setting all simulated values above the 50% quantile to forest and all other values to non-forest (our reference class). We normalized values of all simulated layers to a range between zero and one and replicated the simulation of each layer 100 times per autocorrelation scenario, thus producing 300 unique landscape configurations.

**Figure 4:**
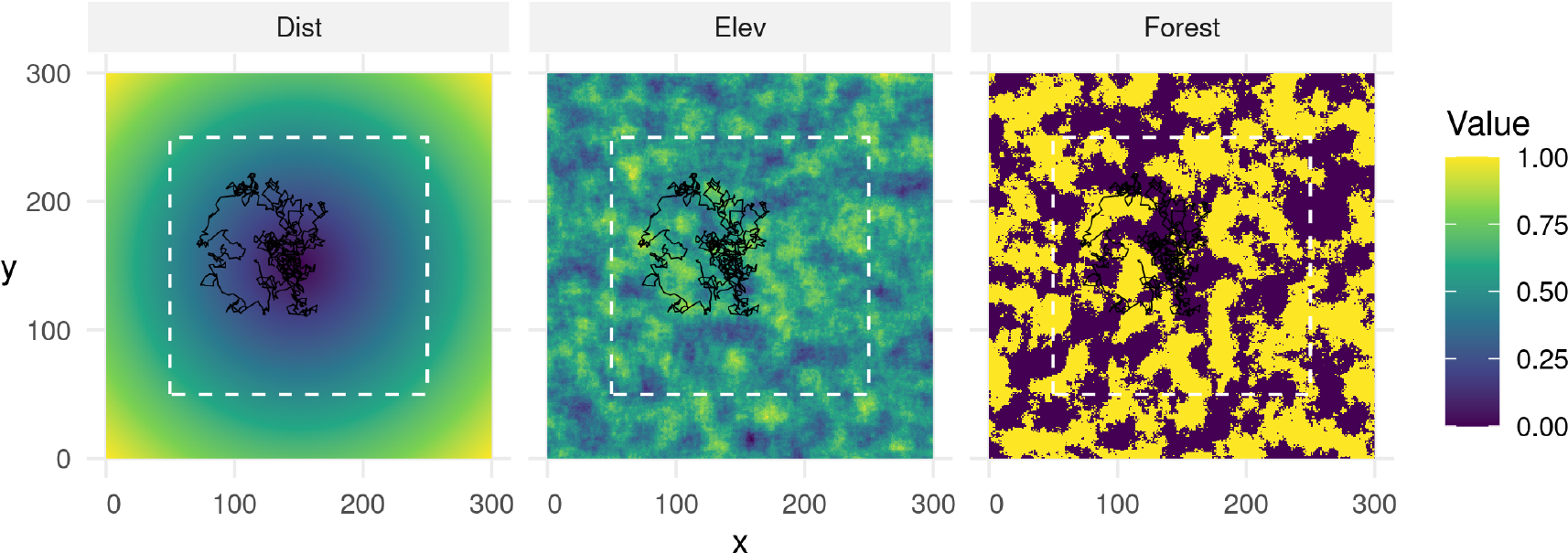
Example of a landscape configuration across which we simulated movement trajectories. All simulated layers had a resolution of 300 x 300 pixels. The distance layer indicated the distance to the center of the landscape and served to simulate attraction. The elevation and forest layers were simulated by sampling pixel-values from a normal distribution and applying a moving window to achieve spatial autocorrelation. Simulated individuals were initiated within the white dashed rectangle, which ensured that they would not be released directly at a map border. Simulated individuals were attracted to the landscape’s center, preferred elevated areas, and avoided areas covered by forest. The black line shows the simulated trajectory associated with the visualized landscape configuration (cfr. Section 2.2).

### 2.2 Movement Simulation

To simulate movement across the virtual landscape, we employed the iSSF simulation algorithm developed by Signer et al. (2017) and applied in Hofmann, Cozzi, McNutt, et al. (2023). This procedure consists of a sequence of five steps that are repeated *n* times to generate a movement trajectory. In step one, we generated a random starting location by sampling random x- and y-coordinates on the simulated landscape. To prevent starting points near map borders, we restricted sampled locations to x- and y-coordinates between 50 and 250 (white dotted rectangle in Figure 4). In step two, we generated a set of 10 random steps originating from the current location, by sampling turning-angles from a von Mises distribution with concentration parameter *κ* = 0.5 and step-lengths from a gamma distribution with shape parameter *k* = 3 and scale parameter *θ* = 1. In step three, we computed average covariate values along each random step from the underlying covariate layers. In step four, we assigned to each step *j* a probability *π*_*j*_ of being selected using the equation below:

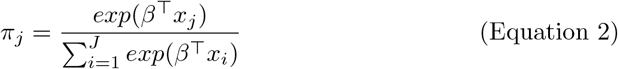

Here, *β* represents the vector of habitat-selection parameters and *x*_*j*_ average covariate values along the *j*^th^ step. The probability of a step being selected thus depended on its associated covariate values, the covariate values of all other random steps, and the simulated preferences *β*. We defined the habitat-selection parameters as *β*_*dist*_ = −20, *β*_*elev*_ = 0.5, and *β*_*forest*_ = −1. That is, simulated individuals were attracted to the landscape’s center, preferred elevated areas, and avoided areas covered by forest. In step five, we sampled one of the random steps based on predicted probabilities and computed the simulated individual’s new position. We then repeated steps two through five until the trajectory comprised a total of 1,000 steps. Each simulated step was assumed to have a step-duration of exactly Δ*t* = 1. We repeated the simulation for each of the 300 simulated landscapes, producing 300 unique trajectories (example trajectory presented in Figure 4).

### 2.3 Data Rarefication

To simulate missingness, we rarefied the trajectories by randomly removing a fixed fraction of animal locations. To assess the impact of different degrees of missingness, we varied the fraction of removed data from 0% (complete dataset) to 50% in increments of 10%. The random removal of animal locations introduced temporal irregularity, such that the resulting step-durations differed depending on the time elapsed between remaining fixes. We replicated the rarefication of each trajectory 100 times.

### 2.4 Computing Bursts

We used the rarefied data to compute bursts of data consisting of a sequence of animal locations with step-durations that did not exceed the forgiveness value. To test how different levels of forgiveness impacted our results, we varied forgiveness from 1 (maximum allowed step-duration was Δ*t* = 1) to 5 (maximum allowed step-duration was Δ*t* = 5). As an example, if the forgiveness was 1, any step with step-duration Δ*t >* 1 resulted in a new burst. If the forgiveness was 2, in contrast, step-durations of up to Δ*t* = 2 were allowed before a new burst was introduced (Figure 1b). Within each burst, we calculated step-lengths and turning-angles. However, due to the grouping of steps into bursts, the orientation of the first step within each burst relative to the previous step could not be determined (Figure 1b). As a result, this step always lacked a turning-angle and was considered invalid.

### 2.5 Fitting Distributions

Based on the steps retained within bursts, we parametrized tentative step-length and turning-angle distributions. Specifically, we used the `fit_distr` function from the amt package (Signer et al., 2019), which is a wrapper function for the `fitdist` function from the `fitdistrplus` package (Delignette-Muller & Dutang, 2015), and fitted a gamma distribution to step-lengths and a von Mises distribution to turning-angles. Notably, we employed two different fitting procedures:

#### 1. Regular Distributions

In this procedure, we fitted parametric distributions considering only step-lengths and turning-angles from steps that exhibited a step-duration of Δ*t* = 1 (i.e., the regular step-duration). Any steps with irregular step-durations (Δ*t >* 1) were discared and did not affect distributional parameter estimates. This represents the traditional proredcure in iSSFs where only regular bursts of animal locations are considered to estimate tentative movement parameters.

#### 2. Dynamic Distributions

In this procedure, we fitted separate parametric distributions to step-lengths and turning-angles from steps of different step-durations. That is, we parametrized separate turning-angle and step-length distributions representative of steps with durations of Δ*t* = 1, 2, 3, 4 and 5 (which corresponds to the maximum forgiveness level we tested for). Some step-durations only rarely occurred at low levels of missingness, thus complicating parametrization of the associated distributions. To facilitate estimation of dynamic distribution parameters across all Δ*t* (Figure S3), we resampled data to different step-durations using the `track_resample` function from the amt package (Signer et al., 2019) before fitting tentative parameters. This ensured a sufficient number of steps for each step-duration to estimate associated parameters.

### 2.6 Step-Selection Functions

We implemented a baseline *uncorrected* approach and four alternative iSSF approaches that mainly differed in the way in which random steps were generated, but sometimes also in the model call that was used to estimate parameters (Figure 3). In the *uncorrected* approach, we treated data as if they were regular, ignoring potential issues arising from differing step-durations. When forgiveness was one, this approach corresponded the traditional iSSF approach. All other approaches were targeted towards reducing potential biases arising from the inclusion of steps with irregular step-durations. Irrespective of the approach employed, we paired each observed step with a total of 10 random steps:

- *Uncorrected*: In the uncorrected approach, we generated random steps by sampling step-lengths and turning-angles from statistical distributions fitted to steps with step-durations of Δ*t* = 1, regardless of the forgiveness value or observed step-durations. Thus, this approach ignored any potential effect of step-duration when generating random step-lengths and turning-angles.
- *Imputed*: In this approach, we sampled step-lengths and turning-angles from statistical distributions fitted to observed steps with a step-duration of Δ*t* = 1. However, prior to generating random steps, we imputed missing fixes using predictions from a simple movement model. Specifically, we fitted a single-state movement model (Johnson et al., 2008) to the simulated trajectories and used the parametrized model to predict coordinates for all missing animal locations. For this, we used the functions `crwFit` and `crwPredict` from the `crawl R`-package (Johnson et al., 2022). Although the `crwFit` function provides capabilities to incorporate location measurement error, we assumed animal locations were measured without error. The imputation resulted in a complete dataset without any missing animal locations, such that each trajectory consisted of a single continuous burst of locations equally spaced in time.
- *Naïve*: In the *naïve* approach, we again sampled step-lengths and turning-angles from regular distributions fitted to steps with step-durations of Δ*t* = 1. However, we linearly scaled the sampled step-lengths depending on the step-durations of the observed steps. For instance, we doubled the sampled step-lengths for any observed step with a step-duration of Δ*t* = 2. This approach naïvely assumed that step-lengths scale linearly with step-durations, which is unlikely to be true, as most animals don’t move in straight lines between successive observations. Furthermore, the linear approximation is likely to get worse as step-duration increases (i.e., as the forgiveness value increases). Since it is not clear how turning-angles should scale with step-duration, we did not adjust the sampled turning-angles.
- *Dynamic+Model*: In the *dynamic+model* approach, we sampled step-lengths and turning-angles from dynamic distributions that were fit to different step-durations. That is, for observed steps with step-duration of Δ*t* = 2, we sampled step-lengths and turning-angles from distributions fit to observed steps with Δ*t* = 2. We then included interactions between the step-duration and other step-descriptors (e.g., sl, log(sl), cos(ta)), allowing us to update movement parameters for each step-duration separately. To avoid numerical instabilities with the conditional logistic regression model, we only included steps with durations Δ*t >* 1 if the respective duration was represented at least 5 times in the rarefied dataset.
- *Multistep*: In the *multistep* approach, we sampled step-lengths and turning-angles from statistical distributions fitted to observed steps with step-durations of Δ*t* = 1. We then generated a sequence of random steps such that their combined step-duration equaled the step-duration of each observed step. For instance, for an observed step with step-duration of Δ*t* = 2, we generated sets of two random steps, which we then concatenated into a “random path”. The paths were then simplified to straight lines connecting the first and last coordinate of each path, which represented the final random step.

Together, an observed step and its 10 associated random steps formed a *stratum* that received a unique ID. Along all steps, we computed average covariate values from the underlying covariate layers.

### 2.7 Conditional Logistic Regression Model

We estimated simulation parameters for the combinations presented in Figure 3 through conditional logistic regression, implemented using the `clogit` function in the `R`-package `survival` (Therneau et al., 2023). We defined a binary response variable (observed) indicating if a step was an observed (scored 1) or a random step (scored 0) and used the step’s ID as a stratification variable. We included habitat covariates (dist, elev, forest) and step-descriptors (sl, log(sl), cos(ta)) as predictors in the regression model. For the *dynamic+model* approach, we also included interactions between the step-duration, coded as a factor, and step-descriptors. To update tentative movement parameters (denoted by the subscript _0_) and obtain the habitat-selection free movement kernel (denoted by the ^ symbol), we employed the formulas provided in (Avgar et al., 2016; Fieberg et al., 2021):

Updated shape 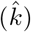 and scale 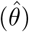 parameters of the step-length distribution (gamma):

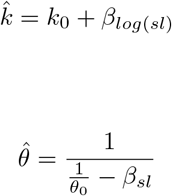

Updated concentration parameter 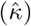 of the turning-angle distribution (von Mises):

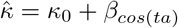

We kept track of the updated movement estimates (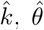, and 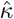), as well as the habitatselection estimates 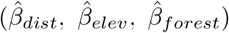, and we quantified bias (estimate−truth) and model accuracy via the root-mean-square error (RMSE).

## 3 Results

All of the approaches were able to recover the parameters of the habitat-selection function, but the *uncorrected, naïve*, and *imputation* approaches resulted in biased estimators of the parameters in the movement kernel, particularly for high forgiveness values (Figure 5). Results were qualitatively similar for all landscape autocorrelation scenarios and for different combinations of missingness and forgiveness (Figure S3). Here, we report on results for a landscape with autocorrelation of 20, while either holding constant missingness at a conservative 20% (Figure 5) or the forgiveness level at 2 (Figure 6) (results for all other combinations are summarized in Figure S3). When missingness was set to 20%, increasing the forgiveness from 1 to 5 improved the precision of the estimators of habitat-selection parameters without introducing bias, irrespective of the chosen method (Figure 5a). Consequently, the RMSE substantially decreased for the habitat-selection parameter estimators as forgiveness increased (Figure 5b). These results highlight the potential benefits of leveraging additional data compared to the traditional approach (represented by the *uncorrected* approach and forgiveness = 1), which uses only bursts of regular data. By contrast, increasing the forgiveness under the *uncorrected, naïve*, and *imputation* approaches resulted in biased estimators of the parameters in the movement kernel (Figure 5a). This bias was more pronounced at higher forgiveness levels, leading to higher RMSE values (Figure 5b). The *imputation* approach appeared to perform particularly poorly at estimating parameters for the turning-angle distribution, yet the magnitude of bias was unrelated to the level of forgiveness and largely owed to missingness *per se* (Figure 6). The *dynamic+model* and *multistep* approaches, on the other hand, performed well, resulting in unbiased or nearly unbiased estimators of parameters of the movement kernel, even with high forgiveness and missingness values (Figure 5 and Figure 6). When forgiveness was fixed to 2, missingness negatively influenced the precision and accuracy of estimates, yet its impact could be dampened using the *dynamic+model* and *multistep* approaches (Figure 6).

**Figure 5:**
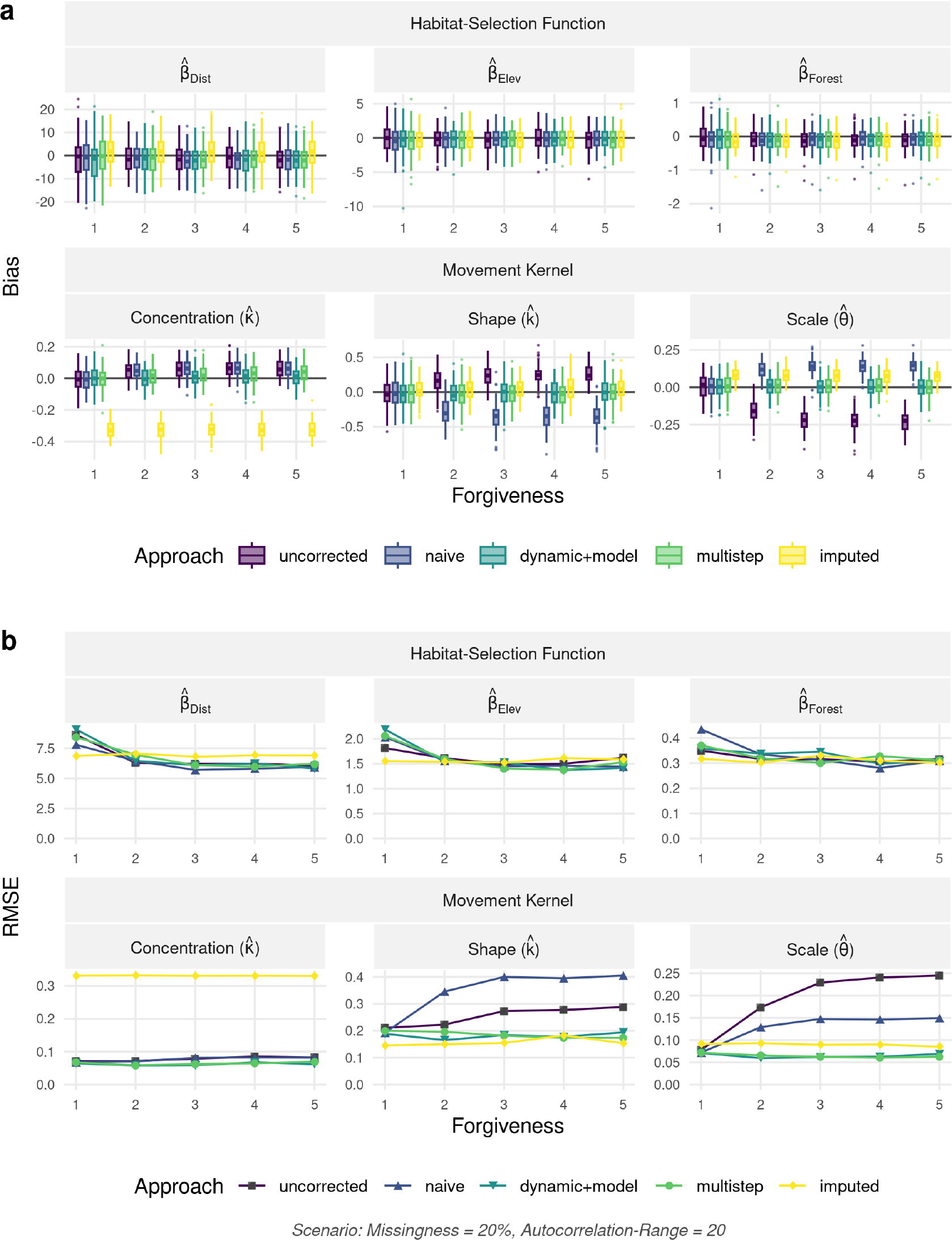
(**a**) Bias (estimate−truth) and (**b**) Root mean-square error (RMSE) with regard to the movement kernel and habitat-selection function as a function of forgiveness. Results are shown for the scenario with landscape autocorrelation of 20 and missingness of 20%. The movement kernel comprised of a gamma distribution with shape parameter *k* and scale parameter *θ* governing the step-length distribution and a von Mises distribution with concentration parameter *κ* governing the turning-angle distribution. Habitat-selection was based on three covariates, namely a Distance, Elevation, and a Forest layer. Estimates are shown for the five different approaches we tested for. The uncorrected approach ignored the fact that higher forgiveness implied temporal irregularity in the data, while all other approaches attempted to correct for the potential biases introduced by temporal irregularity.

**Figure 6:**
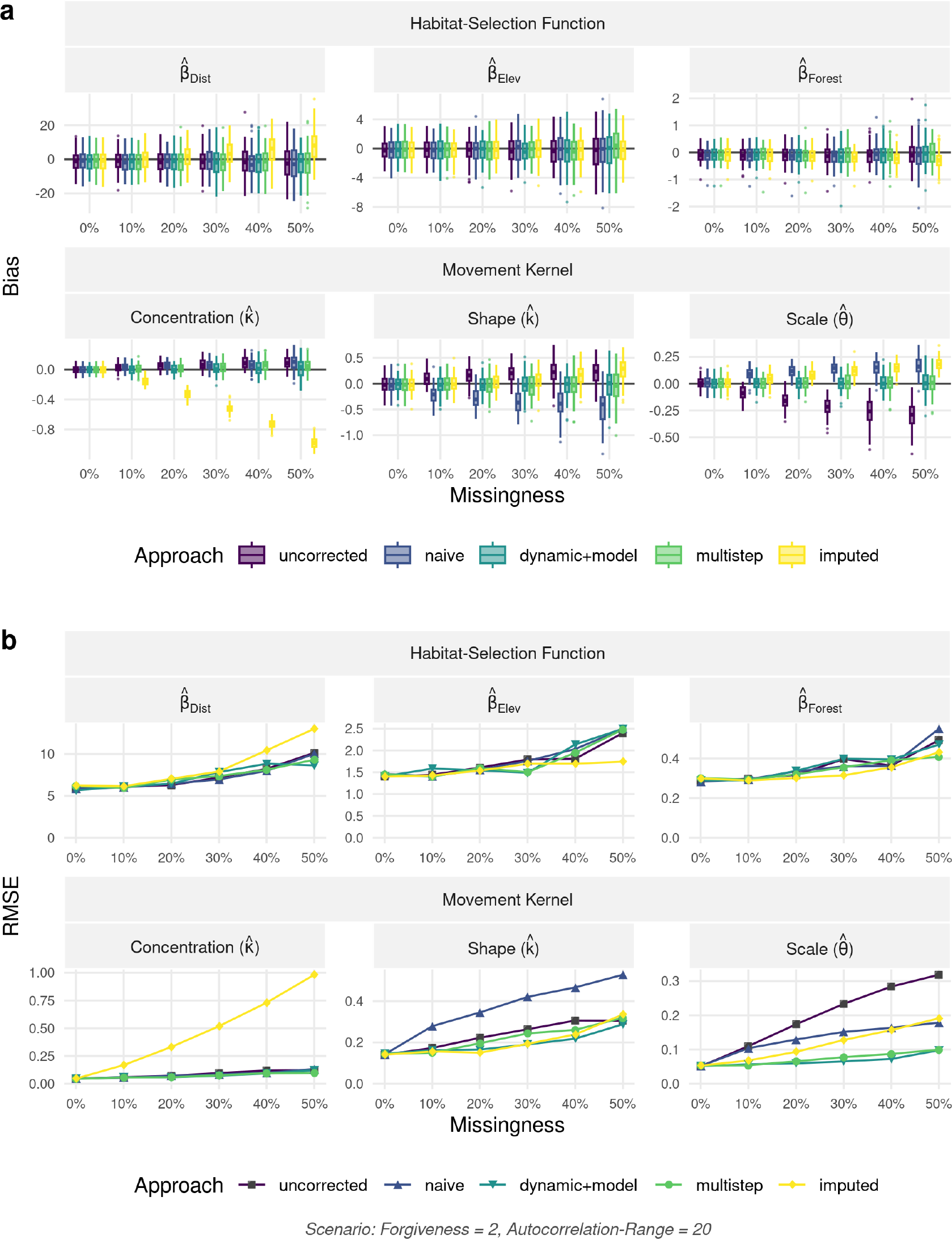
(**a**) Bias (estimate−truth) and (**b**) Root mean-square error (RMSE) with regard to the movement kernel and habitat-selection function as a function of missingness. Results are shown for the scenario with landscape autocorrelation of 20 and forgiveness of 2. The movement kernel comprised of a gamma distribution with shape parameter *k* and scale parameter *θ* governing the step-length distribution and a von Mises distribution with concentration parameter *κ* governing the turning-angle distribution. Habitat-selection was based on three covariates, namely a Distance, Elevation, and a Forest layer. Estimates are shown for the five different approaches we tested for. The uncorrected approach ignored the fact that higher forgiveness implied temporal irregularity in the data, while all other approaches attempted to correct for the potential biases introduced by temporal irregularity.

## 4 Discussion

We conducted a simulation study with known habitat and movement parameters to investigate if retaining irregular animal locations via increased forgiveness improves or worsens parameter estimation in iSSFs. We also tested the performance of four different approaches that attempt to correct for potential biases introduced by using temporally irregular data, and we compared them to an uncorrected baseline approach. Our results demonstrated that retaining irregular animal locations can improve the precision of estimators of habitatselection parameters, but some methods resulted in biased estimators of the parameters in the movement kernel. Overall, our results highlight the potential benefits of leveraging irregular animal locations, especially when an appropriate method for handling irregular data is chosen.

The *uncorrected* baseline approach ignored the fact that increasing forgiveness introduced irregularity in the data. Consequently, estimators of the parameters in the movement kernel were increasingly biased as forgiveness increased due to the inclusion of steps with varying step-durations when fitting the model. Steps with longer step-durations tended to have larger step-lengths and less directed turning-angles, which led to an overestimation of *θ* and underestimation of *κ*. Yet, estimators of the habitat-selection parameters remained unbiased, even at high levels of forgiveness, and they were more precise than the standard estimator (represented by the *uncorrected* approach with forgiveness = 1), highlighting the benefits that can be reaped by including additional data.

Similarly, the *naïve* approach performed well when estimating habitat-selection parameters but resulted in biased estimators of movement parameters, especially for large forgiveness values. This result was unsurprising given that our simulated trajectories were tortuous and therefore step-lengths were not linearly related to step-durations. Indeed, we found that, although there was a near linear relationship between step-duration and the (tentative) scale parameter (*θ*_0_), the relationships between step-duration and the (tentative) movement parameters *κ*_0_ and *k*_0_ were non-linear (Figure S4). Overall, the usefulness of this approach appears highly limited, as it is often not clear by what factor distributional parameters for step-lengths and turning-angles should be multiplied to match the observed step-duration.

The *dynamic+model* approach provided a very flexible, easily implementable, and powerful framework for retrieving precise and unbiased estimates for both habitat-selection and movement parameters, irrespective of forgiveness. Fitting tentative distributions for different step-durations may be challenging due to some step-durations occurring only rarely, yet resampling observed animal locations to different step-durations using the `track_resample` function from the `amt R`-package (Signer et al., 2019) provides an easy solution to generate the needed data. By including interactions between step-descriptors (sl, log(sl), and cos(ta)) and step-duration in the conditional logistic regression model, it was possible to obtain unbiased estimators of movement parameters for steps of varying step-duration. We included step-duration as a categorical covariate, yet there may be times where it would be advantageous to treat it as a continuous covariate (e.g., with its effect modeled using a low-degree polynomial or regression spline with few degrees of freedom). Treating stepduration as a continuous variable may help to alleviate convergence issues in cases where some step-durations are rare, and it might allow applying the *dynamic+model* approach to data that are entirely irregular.

The *multistep* approach also performed well and was relatively easy to implement. This approach is somewhat *ad hoc* in that it uses the tentative movement parameters to generate random steps to match observed steps with longer step-durations (in multiples of Δ*t*). It is similar to, but slightly less principled, than the approach developed by Vales et al. (2022), which formally constructs the likelihood for multistep-durations by integrating out the missing steps. An advantage of this latter approach is that one can also attempt to account for non-random missingness by explicitly modeling factors related to the probability of obtaining a successful location (Vales et al., 2022). Nonetheless, integrating over the missing steps, as in Vales et al. (2022), can be computationally intensive and prohibitive with large data sets. Another downside of both of these approaches (the *multistep* approach and the approach of Vales et al., 2022) is that they can only be applied in cases where step-durations are a fixed multiple of the regular step-duration; i.e., unlike the *dynamic+model* approach, they cannot be applied when data are highly irregular.

Of the methods we considered, the *imputation* approach performed the worst. It resulted in biased estimators of parameters in both the habitat-selection function and the movement kernel, irrespective of the forgiveness value. This bias likely resulted from using an overly simplistic movement model to impute missing fixes. Moreover, the imputation procedure may have led to imputed animal locations that masked important selection properties, therefore leading to inaccurate parameter estimates. While this approach appears to perform well with hidden Markov movement models (McClintock, 2017), we advise against its use with iSSFs.

For the scenarios we considered in our simulation study, the estimators of habitatselection parameters were insensitive to the inclusion of irregular data and performed well, with the exception of the *imputation* approach. This suggests that accounting for irregular step-durations may not be particularly important if one is only interested in the habitat-selection function. When the movement kernel is also of interest, we suggest the *dynamic+model* approach, since it is flexible, easy to implement, and allows one to use more data than the traditional approach that requires bursts of regular data, leading to more precise estimators.

Several authors have emphasized that movement and habitat-selection parameters in an SSF are scale dependent and should be expected to change as the sampling frequency changes (see for example Avgar et al., 2016; Signer et al., 2017; Fieberg et al., 2021). Furthermore, Barnett and Moorcroft (2008) developed an analytical framework for investigating scale dependence and showed that habitat-selection parameters should depend on the relative width of the movement kernel in relation to habitat heterogeneity. Thus, the relative insensitivity of the habitat-selection parameters to the inclusion of steps with varying step duration was somewhat unexpected. It would be interesting to explore the robustness of this result across a wider range of simulation scenarios in the future.

More generally, the spatial scale of a habitat-selection analysis has been recognized as an important factor, which is why Johnson (1980) proposed a hierarchical framework for examining habitat-selection across different orders (e.g., species range, individual home range, within a home range). Johnson’s proposed framework acknowledges that habitat-selection may act differently at different scales and that the interpretation of ecological processes changes depending on the spatial scale at which they are investigated (Wiens, 1989; Levin, 1992). This understanding has encouraged scientists to conduct extensive scaling analyses and to comprehensively examine habitat-selection at multiple scales (DeCesare et al., 2012; McGarigal et al., 2016; Pitman et al., 2017; Zeller et al., 2017). In studies employing SSAs, the issue of scale is often neglected, and data are most frequently analyzed at the spatio-temporal scale at which they were collected. This choice maximizes the number of locations that can be used in the analysis, yet prevents a thorough understanding of scaledependency. The use of irregular data in SSAs poses another challenge, as steps with unequal step-durations may reflect selection processes occurring at different scales. The severity of this issue obviously depends on the original sampling frequency, the degree of missingness, and the scale at which animals are making decisions that are relevant in terms of their movement behavior and habitat selection. By including irregular animal locations via increased forgiveness, we may therefore average over selection processes occurring at multiple scales, which could produce estimates of habitat-selection parameters that are misleading due to contradictory effect-signs at different scales. To better account for such scale-dependent processes, it may be beneficial to include interactions between step-duration and habitat features (e.g., `dist, elev, forest`), thus allowing habitat-selection parameters to also vary as a function of step-duration.

It is important to note that we considered a limited number of scenarios in our simulation study. For instance, we assumed that animal locations were missing at random, i.e., failure to obtain a fix was unrelated to habitat types, time of the day, etc. However, several studies have shown that missingness is often non-random and related to difficulties with satellite transmission due to topography (Lewis et al., 2007), canopy cover (Phillips et al., 1998; DeCesare et al., 2005; Hansen & Riggs, 2008), time of the day (Graves & Waller, 2006), animal behavior (Mattisson et al., 2010), or collar orientation (D’eon & Delparte, 2005). In fact, Vales et al. (2022) highlighted that missingness and the associated under-representation of certain habitat types may lead to biased estimators of parameters in iSSFs, but that accounting for the probability of obtaining a location in differing environmental conditions may alleviate this bias. Future studies should strive to further investigate these relationships and examine how our proposed approaches perform when missingness is habitat-dependent. In addition, future studies could investigate scenarios in which individuals alter their movement tendencies in response to local environmental features (i.e., models with habitat-dependent movement kernels) and examine how this influences the robustness of our proposed approaches.

While our results suggest that irregularity due to missing animal locations can effectively be accounted for in iSSFs and that increasing the forgiveness, thus allowing for inclusion of irregular data, improves estimator precision, we also found a decreasing marginal benefit of increased forgiveness. In fact, increasing the forgiveness beyond a value of two (i.e., allowing for steps of twice the regular step-duration) only marginally improved model performance in our case. This can also be seen in Figure 2, which shows that the largest number of steps that can be gained is when increasing the forgiveness level from 1 to 2. Having a higher forgiveness beyond 2 may thus not even be necessary, therefore limiting the need to correct biases emerging from the inclusion of irregular data.

Our study contributes to the growing body of literature that extends iSSFs and improves the method’s robustness under various conditions. This includes approaches for modeling irregular data (Munden et al., 2021), accounting for spatial dependence among residuals (Arce Guillen et al., 2023), methodological frameworks for fitting iSSFs with random slopes (Muff et al., 2020), as well as incorporating the probability of successfully obtaining an animal location in different habitat conditions (Vales et al., 2022).

In conclusion, our study shows that inclusion of irregular animal locations can improve model performance, yet only when an appropriate approach to account for irregularity is selected. Here, the *dynamic+model* and *multistep* approaches performed well and resulted in improved estimators of habitat-selection and movement parameters, even at elevated levels of missingness and forgiveness. Both methods are easy to implement, and the associated models can readily be fitted using conditional logistic regression analysis, such as provided through the R-packages `amt` (Signer et al., 2019), `survival` (Therneau et al., 2023), or coxme (Therneau, 2022). To facilitate uptake and encourage use of the proposed approaches among practitioners, we provide all of our codes through an online repository, which includes an example application of the *dynamic+model* approach. With this, we hope practitioners will rethink the common use of discarding large portions of data and instead use methods that can appropriate accomodate irregular data.

## 5 Authors’ Contributions

D.D.H., G.C., and J.F. conceived the study and designed methodology; D.D.H. implemented the analysis, J.F. assisted with modeling; D.D.H. wrote the first draft of the manuscript and all authors contributed to the drafts at several stages and gave final approval for publication.

## 6 Data Availability

Code to reproduce this study is available through the University of Minnesota’s Data repository (https://hdl.handle.net/11299/257804, Hofmann, Cozzi, and Fieberg, 2023). The repository also includes an example analysis that showcases the application of the *dynamic+model* approach.

## 7 Acknowledgements & Funding

D.D.H. was funded by a Swiss National Science Foundation Grant No. 310030 204478 rewarded to G.C. J.F. was supported by the National Aeronautics and Space Administration award 80NSSC21K1182 and received partial salary support from the Minnesota Agricultural Experimental Station.

