## Supporting Information for "Methods for Implementing Integrated Step-Selection Functions with Incomplete Data"

November 8, 2023

[1] Department of Evolutionary Biology and Environmental Studies, University of Zurich, Winterthurerstrasse 190, 8057 Zurich, Switzerland.

[2] Botswana Predator Conservation Program, Wild Entrust, Private Bag 13, Maun, Botswana.

[3] Department of Fisheries, Wildlife, and Conservation Biology, University of Minnesota, St. Paul, MN, USA.

**Running Title:** Step-Selection Analyses with Missing Data

**Keywords:** animal movement, gps data, imputation, incomplete data, missing fixes, step-selection analyses, step-selection functions

### A.1 Landscape Simulation: Different Autocorrelation Scenarios

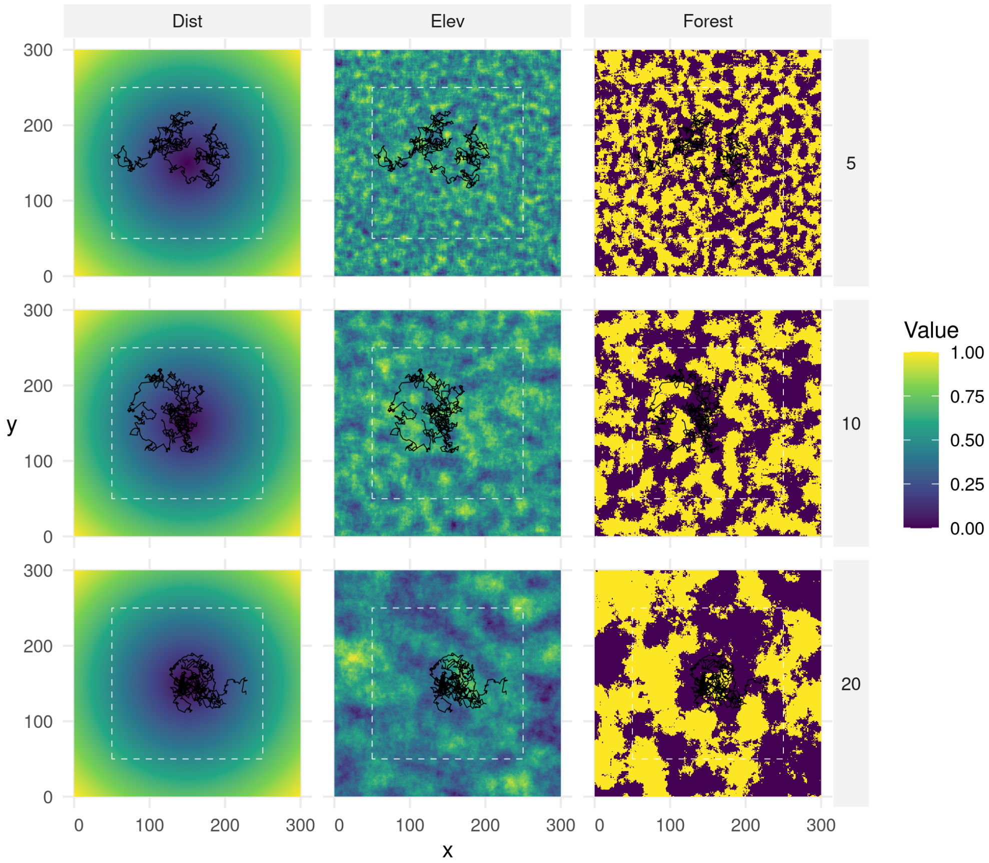


**Figure S1:** Simulated landscapes under different levels of autocorrelation (5, 10, 20; from top to bottom). Autocorrelation only affected the layers elev and forest, which were both simulated using a Gaussian random field neutral landscape model (Schlather et al., 2015) using the R-package `NLMR` (Sciaini et al., 2018). Simulations were repeated 100 times for each autocorrelation scenario, thus resulting in 300 unique landscape configurations.

### A.2 Dynamic Tentative Distribution Parameters

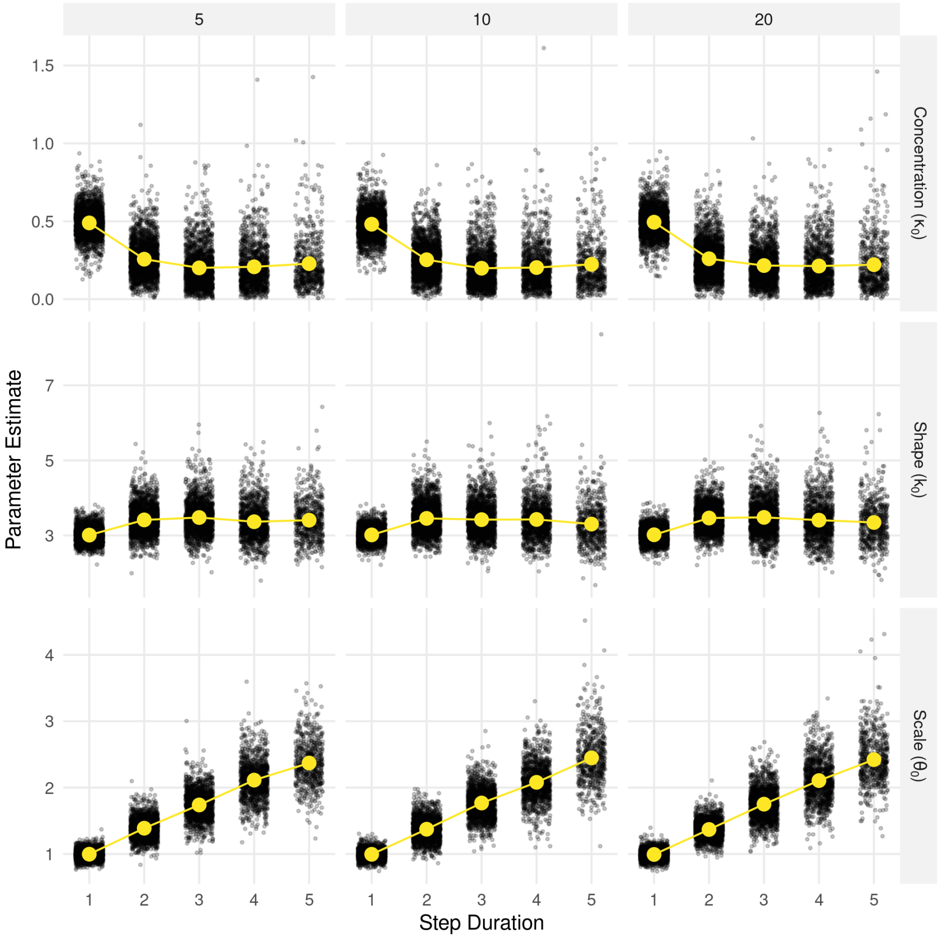


**Figure S2:** Tentative prameter estimates for the von Mises distribution (top row) and gamma distribution (bottom row) fitted to steps with different durations. The von Mises distribution requires one parameter, namely a concentration parameter ($\kappa$). The gamma distribution requires a shape parameter ($k$) and a scale parameter ($\theta$). The subscript $_0$ is used to indicate that these are tentative distribution parameters (sensu Avgar et al., 2016 and Fieberg et al., 2021).

### A.3 Model Estimates across all Scenarios

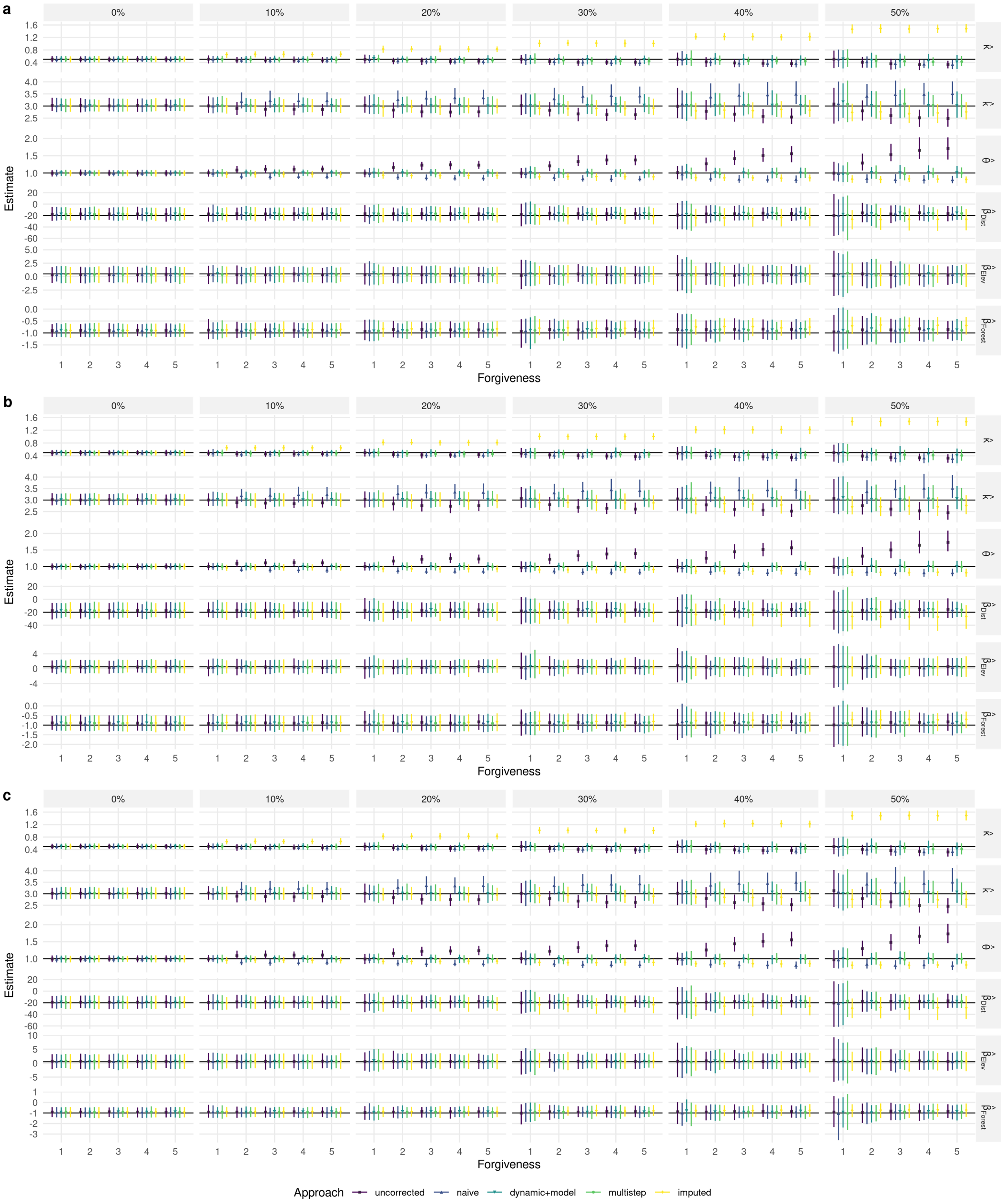


**Figure S3:** Parameter estimates across different autocorrelation scenarios (5, 10, 20; panels a, b, and c) and missingness levels (0% - 50%; from left to right). True simulation parameters are indicated by the solid black lines. Parameter estimates from the different approaches are given by the colored symbols, and their bootstrap 95% CIs across 100 replicates by the colored lines.
